# Cardiac remodeling after six weeks of high-intensity interval training to exhaustion in endurance-trained males

**DOI:** 10.1101/582924

**Authors:** Haïfa Mahjoub, Olivier Le Blanc, Myriam Paquette, Sarah Imhoff, Lawrence Labrecque, Audrey Drapeau, Paul Poirier, Élisabeth Bédard, Philippe Pibarot, Patrice Brassard

**Affiliations:** Department of Medicine, Université Laval, Québec, Canada; Department of Kinesiology, Faculty of Medicine, Université Laval, Québec, Canada; Research center of the Institut universitaire de cardiologie et de pneumologie de Québec, Québec, Canada; Faculty of Pharmacy, Université Laval, Québec, Canada

**Keywords:** high-intensity interval training, endurance-trained individuals, ambulatory blood pressure, left atrium, right ventricle

## Abstract

High-intensity interval training (HIIT) improves physical performance of endurance athletes, although studies examining its cardiovascular effects are sparse. We evaluated the impact of HIIT on blood pressure, heart rate and cardiac cavities size and function in endurance-trained adults. Seventeen endurance-trained males underwent 24-hour ambulatory blood pressure monitoring (ABPM) and Doppler echocardiography at baseline and after 6 weeks of HIIT. Participants were divided in 2 groups (85% maximal aerobic power (MP); HIIT_85_, n=8 and 115% MP; HIIT_115_, n = 9) in order to compare the impact of different HIIT intensities. ABPM and cardiac chambers size and function were similar between groups at baseline. HIIT reduced heart rate (55 ± 8 vs. 51 ± 7 bpm; p= 0.003), systolic blood pressure (SBP; 121 ± 11 vs 118 ± 9 mmHg; p = 0.01), mean arterial pressure (MAP; 90 ± 8 vs 89 ± 6 mmHg; p = 0.03) and pulse pressure (52 ± 6 vs 49 ± 5 mmHg; p = 0.01), irrespective of training intensity. Left atrium (LA) volumes increased after HIIT (maximal: 50 ± 14 vs 54 ±14 ml; p= 0.02; minimal: 15 ± 5 vs 20 ± 8 ml; p = 0.01) in both groups. Right ventricle (RV) global longitudinal strain lowered after training in the HIIT_85_ group only (20 ± 4 vs. 17 ± 3%, p = 0.04). In endurance-trained males, six weeks of HIIT reduce SBP and MAP and increase LA volumes irrespective of training intensity, whereas submaximal HIIT deteriorates RV systolic function.

**Key points:**

- High-intensity interval training improves physical performance of endurance athletes, although studies examining its cardiovascular effects are sparse.
- We evaluated the impact of submaximal (85% maximal aerobic power) and supramaximal (115% maximal aerobic power) high-intensity interval training on ambulatory blood pressure, heart rate and cardiac cavities size and function in endurance-trained adults.
- Irrespective of training intensity, six weeks of high-intensity interval training increase left atrial volumes in endurance-trained adults, whereas the submaximal training decreases right ventricular systolic function.
- These results may help identify the exercise threshold for potential toxicity of intense exercise training for at-risk individuals and ideal exercise training regimens conferring optimal cardiovascular protection and adapted endurance training for athletes.

## INTRODUCTION

High-intensity interval training (HIIT) involves repeated intervals of high-intensity aerobic activity interspersed with rest or active lower-intensity recovery periods. There is growing interest in HIIT because of robust and ever-increasing evidence of its efficacy for improving cardiovascular and metabolic functions in both healthy populations and populations with a chronic disease (Ramos *et al*., 2015; Batacan *et al*., 2017). Owing to its capacity to improve maximal oxygen consumption (VO_2_ max) and physical performance, HIIT is already a main part of endurance athlete training (Buchheit & Laursen, 2013). However, little is known about the impacts of this training modality on cardiac haemodynamics and cardiac remodeling in endurance-trained individuals.

Exercise training induces a decrease in blood pressure (BP) in healthy untrained (Cornelissen & Smart, 2013) or hypertensive individuals (Ciolac *et al*., 2009). Nonetheless, the impact of training intensity on BP in athletes remains unclear. HIIT appears to influence favorably BP response to exercise but not resting or ambulatory BP (Berge *et al*., 2015).

Otherwise nearly half of aerobically-trained individuals present some evidence of cardiac remodeling, which consists of changes in chamber dimensions including increased left and right ventricular and atrial cavity size associated with normal systolic and diastolic function (Maron & Pelliccia, 2006). These structural alterations represent cardiac adaptations to the high haemodynamic demands of exercise and have been considered typical findings of the athlete’s heart (Pluim *et al*., 2000; Maron & Pelliccia, 2006); they can be observed after only few months of intensive training (D’Ascenzi *et al*., 2012) and can be reversed after a detraining period (D’Ascenzi *et al*., 2015). However, accumulating data suggest that training of elite endurance athletes including those competing in extreme endurance events (i.e. marathons, ultramarathons, ironman distance triathlons) can cause transient acute volume overload of cavities, that may lead after months to years of repetitive injury to myocardial fibrosis creating a substrate for atrial and ventricular arrhythmias (O’Keefe *et al*., 2012).

The purpose of this study was to evaluate the effects of HIIT on cardiac haemodynamics, (i.e. ambulatory BP and heart rate) and on cardiac cavities size and function in endurance-trained individuals. We also compared submaximal and supramaximal HIIT protocols, which represent different training methods commonly used by endurance athletes to increase VO_2_max (Westgarth-Taylor *et al*., 1997; Weston *et al*., 2014). We hypothesized that HIIT would decrease ambulatory BP and heart rate, but enlarge cardiac cavities without influencing negatively cardiac function.

## METHODS

### Ethical approval

The local ethics committee approved this study (CER:20869) according to the principles established in the Declaration of Helsinki (except for registration in a database), and all participants provided written informed consent.

### Participants

Nineteen endurance-trained males with a training history of 5 to 12 hrs/week for at least 2 years, volunteered to participate in this study. Participants were practicing different types of endurance sports; cycling (*n* = 9), triathlon (*n* = 7), mountain biking (*n* = 2) and cross-country skiing (*n* = 1), and were training 8.4 ± 2.7 hrs/week taking part in 0 to 2 interval training sessions/week (0.5 ± 0.7 HIIT sessions/week) in the month before the study. Although this study was part of a previously published study (Paquette *et al*., 2017) examining HIIT training effects on performance and physiological function in endurance-trained participants, the experimental question addressed in the current paper was determined *a priori* and are exclusively dealt with here.

### Experimental Design

Participants reported to the laboratory on three occasions over two weeks to complete: 1) anthropometrics measurements and Doppler echocardiography at rest, 2) a progressive ramp exercise cycling protocol aiming to determine VO_2_max and installation of the 24-hour ambulatory BP monitoring device, and 3) a maximal aerobic power stepwise intermittent protocol on a cycling ergometer to prescribe training intensity. Participants were asked to refrain from training for at least 12 hrs and to avoid alcohol and caffeine consumption for 24 hrs before each visit. After these preliminary evaluations, participants were matched according to their age and VO_2_max before one from each pair was randomly assigned to one of two training groups: 1) submaximal (HIIT_85_) or, 2) supramaximal (HIIT_115_). 24-hour ABPM, Doppler echocardiography at rest and ramp exercise protocol were repeated 48 to 96 hrs following the end of the 6-week training program.

### Training interventions

The complete training interventions have been described previously (Paquette *et al*., 2017). Briefly, the training period was 6-week long, with 3 HIIT sessions/week to exhaustion. Training sessions were separated by 48 to 72 hrs. The HIIT_85_ group performed repeated effort bouts of 1- to 7-min at 85% maximal aerobic power, interspersed by half the effort time of active recovery (150 W or 50% maximal aerobic power if maximal aerobic power < 300 W). The HIIT_115_ group performed repeated effort bouts of 30-s to 1-min at 115% maximal aerobic power, interspersed by twice the effort time of active recovery (150 W or 50% maximal aerobic power if maximal aerobic power < 300 W). Participants from both groups were asked to exercise until exhaustion, defined as the inability to complete an effort bout. Session rate of perceived exertion (0-10 scale) was obtained within 10 min after the end of each training session (Foster, 1998). Exercise bout duration was alternated from one session to another in both groups to reduce exercise training monotony and to focus on exercise intensity (85 vs 115% maximal aerobic power) instead of exercise duration (Paquette *et al*., 2017). HIIT_85_ and HIIT_115_ protocols were matched for total effort rather than for total work, which is closer to what athletes typically do when performing hard interval sessions at various intensities (Paquette *et al*., 2017). On remaining days, participants were asked to avoid HIIT, but to maintain a similarly low and/or moderate intensity training volume as before the study. Training sessions were performed on Tacx Bushido trainers (Tacx, Terneuzen, The Netherlands) and participants utilized their own bike for all training sessions.

### Methodology

Height and body mass were measured in each participant. Maximal aerobic power was measured for the determination of training intensities (85 and 115% maximal aerobic power) as previously described (Paquette *et al*., 2017).

#### Maximal oxygen consumption

VO_2_max was determined using an electromagnetically braked upright cycling ergometer (Corival, Lode, Groningen, The Netherlands). VO_2_, carbon dioxide production (VCO_2_), and respiratory exchange ratio (RER: VCO_2_/VO_2_) were recorded continuously by a breath-by-breath gas analyser (BreezeSuite, MedGraphics Corp., Saint Paul, MN, USA). Following a 3- min of seated rest on the upright cycling ergometer, and 1 min of unloaded pedalling, the ramp incremental protocol started, with 25-30 W/min increments until volitional exhaustion. VO_2_max was defined as the highest 30 s averaged VO_2_ concurrent with a RER ≥ 1.15 (Paquette *et al*., 2017).

#### Ambulatory blood pressure

BP was recorded for 24 hrs (Mobil-O-Graph, model no. B09364, France) as participants were asked to continue their regular daily activities with no more than a moderate intensity. BP was measured every 30 min during the day (6 AM to 11 PM) and every 60 min during the night (11 PM to 6 AM). Systolic blood pressure (SBP), diastolic blood pressure (DBP), mean arterial pressure (MAP) and pulse pressure (PP) were obtained from ambulatory BP monitoring. Resting systemic arterial hypertension was defined as SBP > 135 mm Hg and/or DBP > 85 mm Hg averaged during the 24-hr ambulatory BP monitoring (Chobanian, 2007).

#### Echocardiography

At entry of the study, all participants underwent a comprehensive Doppler echocardiographic evaluation at rest (Philips IE33, Bothell, WA, USA), and underwent a second echocardiography after the 6-week training program. All echocardiographic studies were reviewed by the same cardiologist (HM) blinded to group allocation. Left ventricle (LV), right ventricle (RV), atrial dimensions and function parameters were measured according to the methods recommended by the American Society of Echocardiography (ASE) guidelines (Lang *et al*., 2015). Left atrial (LA) volumes were calculated from apical 4- and apical 2-chamber views of the LA using the biplane method of disks, as recommended by the ASE. LA passive volumes consist of preatrial contraction volume (V_preA_) measured at the onset of the P-wave on an electrocardiogram; minimal LA volume (V_min_) measured at the closure of the mitral valve in end-diastole and maximal LA volume (V_max_) measured just before the opening of the mitral valve in end-systole. Indexes of atrial compliance were calculated as follow: LA expansion index = (V_max_ - V_min_)/V_max_, LA passive emptying fraction (V_max_ - V_preA_)/ V_max_ and LA passive emptying fraction (V _pre A_ - V_min_)/ V_preA_ (Ancona *et al*., 2014). LV ejection fraction (LVEF) was measured by biplane Simpson method. Mitral flow velocity (E wave) was assessed by pulsed-wave Doppler from the apical 4-chamber view by placing a sample volume at the tips of the mitral leaflets in diastole and recording at a sweep speed of 100 mm/s. The average of lateral and medial mitral annulus early diastolic velocity (e’ wave) was measured using tissue Doppler imaging (TDI). The E/e’ ratio was used to estimate filling pressures. LV, RV and LA strain parameters were retrospectively measured using dedicated commercial software for speckle tracking imaging (2D Cardiac Performance Analysis, Tomtec Imaging Systems, Munich, Germany). LV longitudinal strain was measured from the 4-chamber, 2-chamber and 3-chamber apical views, longitudinal strain of the RV free wall (RVLS) and global longitudinal strain of the RV (RVGLS) were measured from the 4-chamber view. RVLS was the average value of the three segments of the RV free wall and RVGLS the average value of the three segments of the RV free wall and the three segments of the interventricular septal wall. LA strain was measured from the 4-chamber and the 2-chamber views, the software generates the longitudinal strain curves for each segment and a mean curve of all segments: peak atrial longitudinal strain (PALS) was measured at the end of the reservoir phase (positive peak during LV systole), peak atrial contraction strain (PACS) just before the start of the active contractile phase (positive peak during early diastole), and negative peak atrial strain (negative PAS) was measured at the end of atrial contraction (Figure 1). LA stiffness was calculated as LASt = (E/e’) /PALS (Machino-Ohtsuka *et al*., 2011).

**Figure 1.**
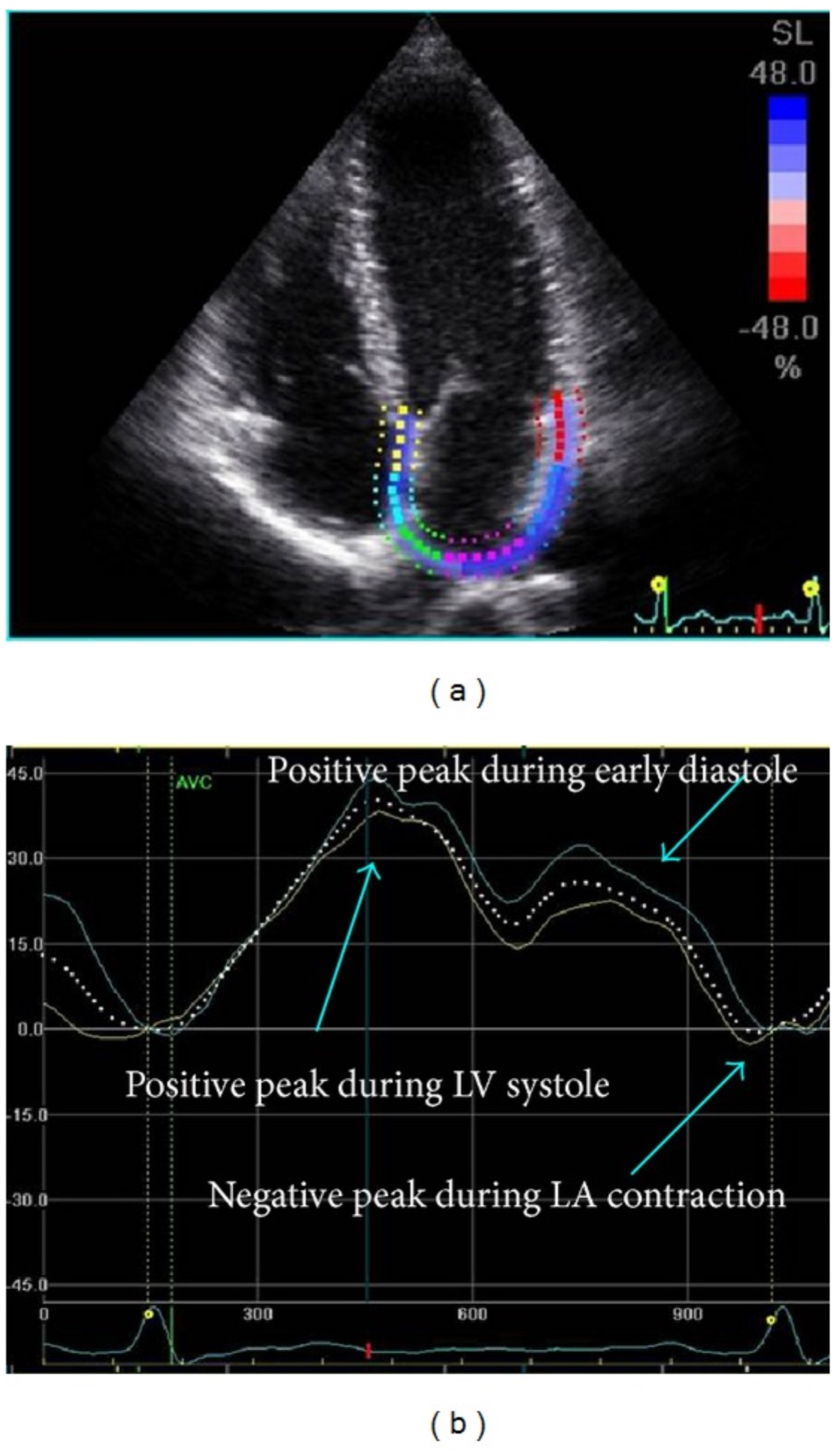
Left atrial strain by speckle tracking. During reservoir function, atrial strain increases, reaching a positive peak (systolic peak), at the end of atrial filling, before the mitral valve opening, during LV systole. After mitral valve opening, during conduit phase, atrial strain decreases, until reaching a plateau, during diastasis, followed by a second positive peak, during early diastole, before LA contraction. Then there is a negative peak, at the end of atrial contraction (Ancona *et al*., 2014).

#### Statistical analysis

Statistical analyzes were performed using SPSS statistical software, version 19.0 (Statistical Package for Social Science, IL, USA). Continuous data were expressed as mean ± standard deviation (SD) unless specified otherwise. After confirmation of distribution normality using Shapiro-Wilk normality tests, between group differences (HIIT_85_ vs. HIIT_115_) were analyzed using a two-way (Training: Baseline vs. Post training x Group: HIIT_85_ vs. HIIT_115_) repeated measures analysis of variance (ANOVA). Following an interaction effect (Training x Group), differences were located using paired samples t-tests, with Bonferroni correction. Relationships were determined using Pearson Product Moment Correlations and statistical significance was established at p < 0.05.

## RESULTS

Seventeen athletes (age: 27 ± 7 years; body weight 72 ± 10 kg) completed the study; 8 participants in the HIIT_85_ group and 9 participants in the HIIT_115_ group. One participant in HIIT_85_ was excluded from analysis due to illness and absence for more than 3 training sessions and one participant in HIIT_115_ was excluded from analysis due to excessive fatigue during the training regime precluding evaluations completion. High-intensity training volume was 47% less in HIIT_115_ group than in HIIT_85_ group (19.3 ± 4.6 vs. 36.5 ± 14.4 min/session; p=0.005). Characteristics of both training intensities, training compliance and averaged rate of perceived exertion have previously been published (Paquette *et al*., 2017). The 6 weeks of HIIT significantly reduced heart rate (55 ± 8 vs. 51 ± 7 bpm; p= 0.003) and improved VO_2_max in all athletes (56 ±5 vs. 59±4 ml/kg/min; p= 0.002) irrespective of training intensity. Body composition measures were not affected by either HIIT program (Paquette *et al*., 2017).

### Impact of HIIT on ambulatory BP

There were no statistically significant differences in ambulatory BP between the HIIT_85_ and HIIT_115_ groups at baseline (Table 1 and Figure 2). Three participants were considered mildly hypertensive at baseline: 2 from the HIIT_85_ group with mean daytime BP of 136/79 mmHg for the first participant and mean daytime BP at 139/77 mmHg for the second participant and one participant from the HIIT_115_ group with mean daytime BP of 141/93 mmHg. After 6 weeks of HIIT, systolic BP, MAP and PP significantly decreased (−5 ± 2 mmHg; p = 0.01; −3 ± 5 mmHg; p = 0.03 and −3 ± 5 mmHg; p = 0.01, respectively), but not DBP. The 3 mildly hypertensive participants normalized their ambulatory BP following HIIT (Table 1 and Figure 2).

**Figure 2.**
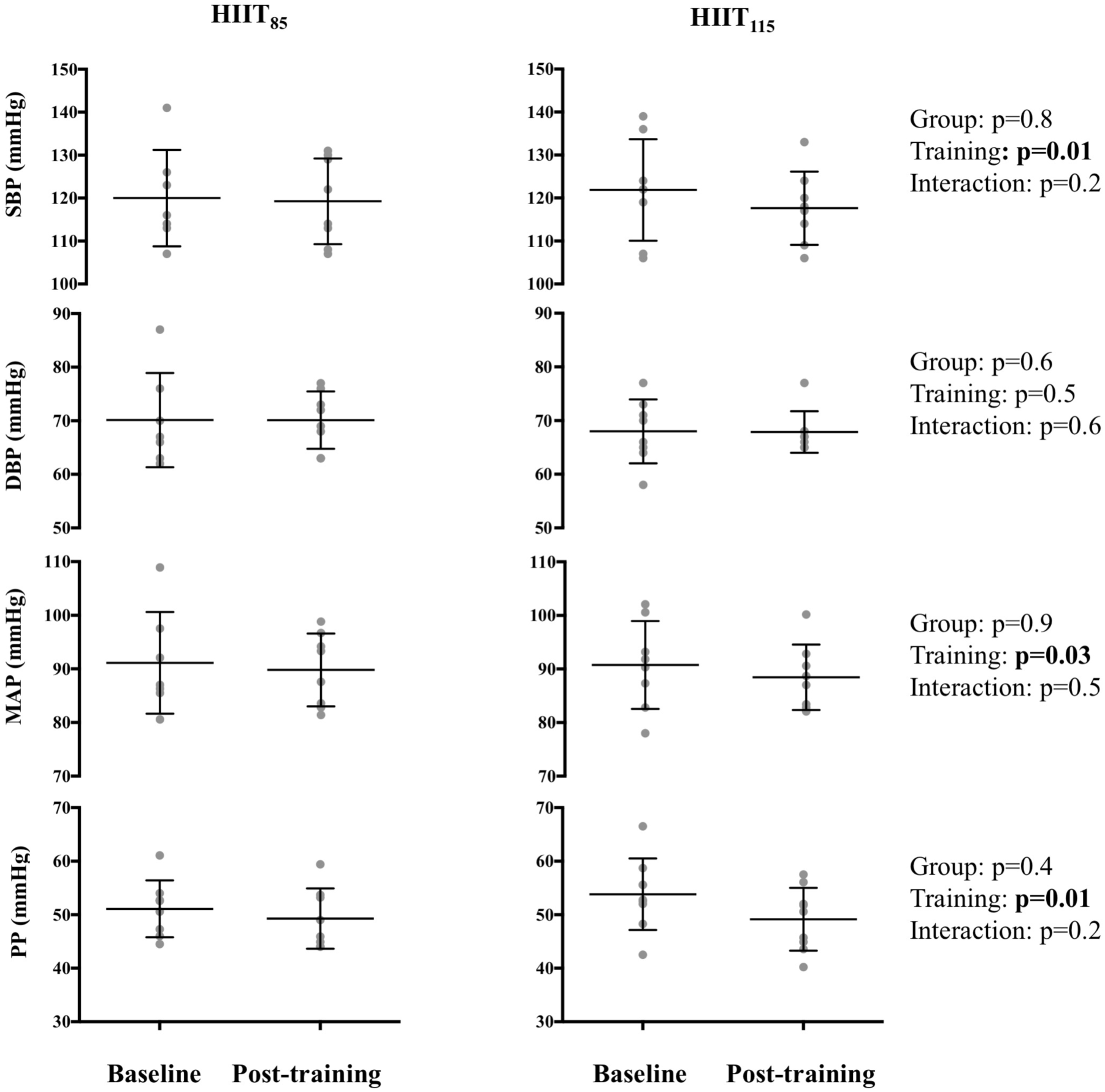
Impact of HIIT on ambulatory blood pressure. DBP: Diastolic blood pressure; HIIT_85_: Submaximal training group; HIIT_115_: Supramaximal training group; MAP: Mean arterial pressure; SBP: Systolic blood pressure; PP: Pulse pressure

**Table 1.** Impact of training on ambulatory blood pressure in endurance athletes from HIIT_85_ and HIIT_115_ groups.

|  | All (n= 17) |  |  | HIIT<sub>85</sub> (n=8) |  |  | HIIT<sub>115</sub> (n= 9) |  |  | p values |  |  |
| --- | --- | --- | --- | --- | --- | --- | --- | --- | --- | --- | --- | --- |
|  | Pre | Post | Delta | Pre | Post | Delta | Pre | Post | Delta | Group | Training | Interaction |
| Systolic blood pressure (mmHg) | 121±11 | 118±9 | **-4.9±2** | 121±11 | 117±8 | -7.4±6 | 120±11 | 119±9 | -2.4±5 | 0.7 | 0.01* | 0.1 |
| Diastolic blood pressure (mmHg) | 69±7 | 69±4 | -0.8±4 | 68±5 | 67±3 | -1.4±3 | 70±8 | 70±5 | -0.2±5 | 0.6 | 0.4 | 0.6 |
| Mean arterial pressure (mmHg) | 90±8 | 89±6 | **-3.3±5.1** | 90 ±8 | 88±6 | -4.4±3 | 91±9 | 89±6 | -2.3±6 | 0.9 | 0.03* | 0.4 |
| Pulse Pressure (mmHg) | 52±6 | 49±5 | **-3.3±4.6** | 53±6 | 49±5 | -4.6±5 | 51±5 | 49±5 | -1.8±3 | 0.6 | 0.01* | 0.2 |

Data are means ± SD

### Impact of HIIT on atria size and function

At baseline, there was no statistical difference in atrial size and function between groups (Table 2). HIIT significantly increased LA volumes; by 19% for maximal LA volumes and by 34% for minimal LA volumes (Figure 3). HIIT was not associated with any change in RA size. Moreover, no significant change was reported in LA function parameters, i.e. indexes of LA compliance and strain parameters (Table 2).

**Figure 3.**
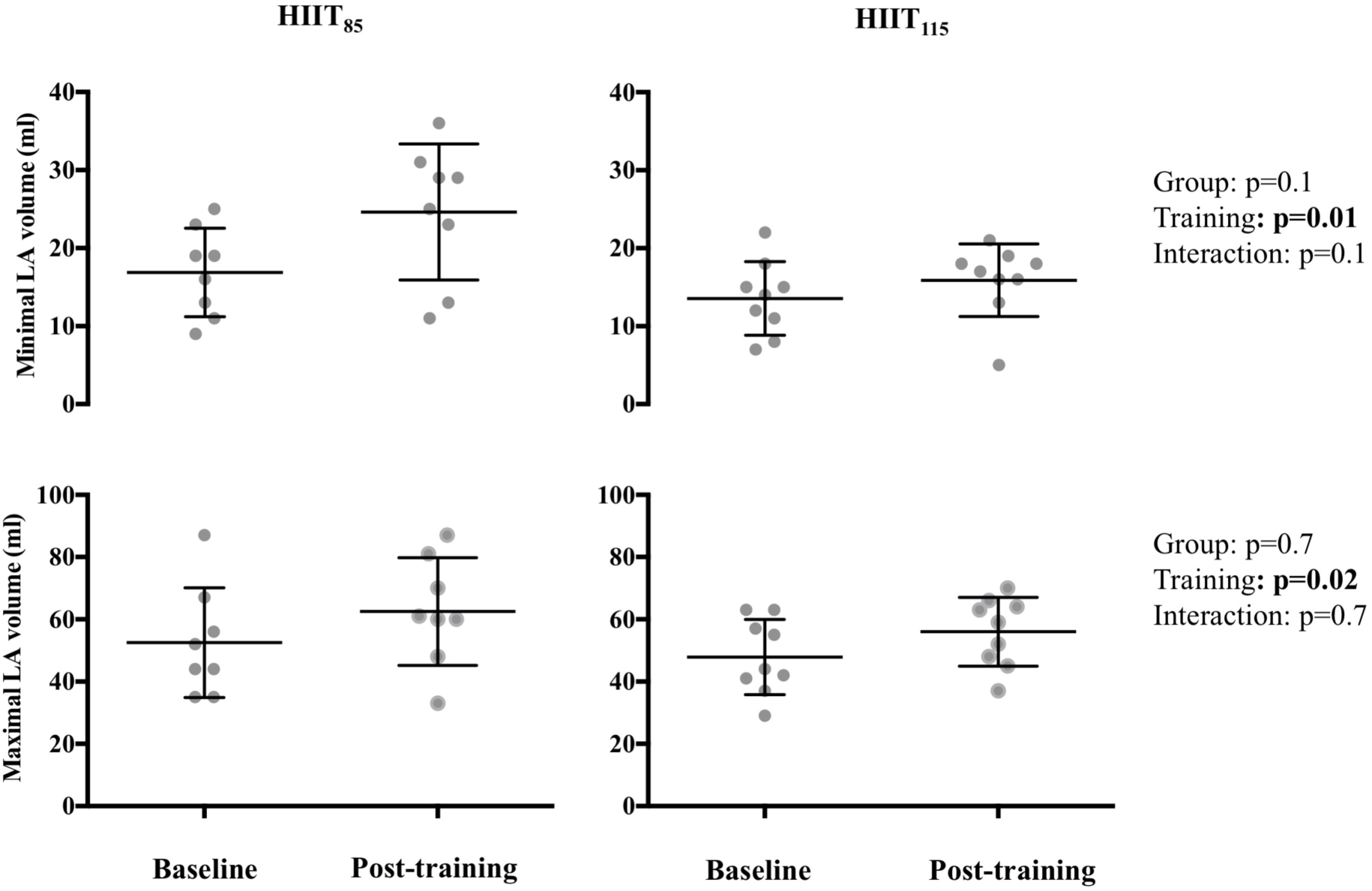
Impact of HIIT on left atrial minimal and maximal volumes. HIIT_85_: Submaximal training group; HIIT_115_: Supramaximal training group; LA: left atrium

**Table 2.**
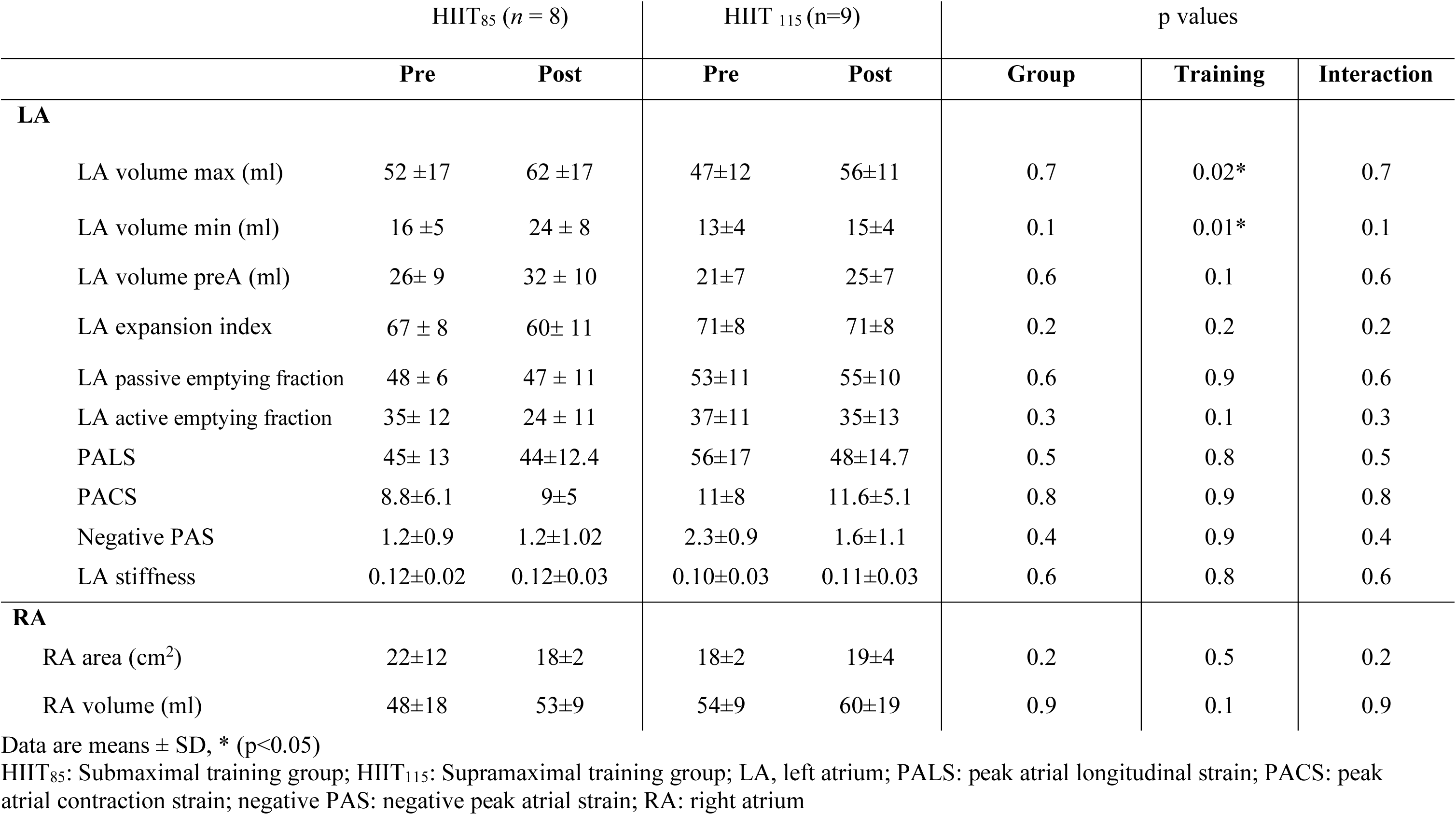
Impact of training on LA and RA size and function in endurance athletes from HIIT_85_ and HIIT_115_ groups.

|  | HIIT<sub>85</sub> (n = 8) |  | HIIT<sub>115</sub> (n=9) |  | p values |  |  |
| --- | --- | --- | --- | --- | --- | --- | --- |
|  | Pre | Post | Pre | Post | Group | Training | Interaction |
| **LA** |  |  |  |  |  |  |  |
| LA volume max (ml) | 52 ±17 | 62 ±17 | 47±12 | 56±11 | 0.7 | 0.02* | 0.7 |
| LA volume min (ml) | 16 ±5 | 24 ± 8 | 13±4 | 15±4 | 0.1 | 0.01* | 0.1 |
| LA volume preA (ml) | 26± 9 | 32 ± 10 | 21±7 | 25±7 | 0.6 | 0.1 | 0.6 |
| LA expansion index | 67 ± 8 | 60± 11 | 71±8 | 71±8 | 0.2 | 0.2 | 0.2 |
| LA passive emptying fraction | 48 ± 6 | 47 ± 11 | 53±11 | 55±10 | 0.6 | 0.9 | 0.6 |
| LA active emptying fraction | 35± 12 | 24 ± 11 | 37±11 | 35±13 | 0.3 | 0.1 | 0.3 |
| PALS | 45± 13 | 44±12.4 | 56±17 | 48±14.7 | 0.5 | 0.8 | 0.5 |
| PACS | 8.8±6.1 | 9±5 | 11±8 | 11.6±5.1 | 0.8 | 0.9 | 0.8 |
| Negative PAS | 1.2±0.9 | 1.2±1.02 | 2.3±0.9 | 1.6±1.1 | 0.4 | 0.9 | 0.4 |
| LA stiffness | 0.12±0.02 | 0.12±0.03 | 0.10±0.03 | 0.11±0.03 | 0.6 | 0.8 | 0.6 |
| **RA** |  |  |  |  |  |  |  |
| RA area (cm<sup>2</sup>) | 22±12 | 18±2 | 18±2 | 19±4 | 0.2 | 0.5 | 0.2 |
| RA volume (ml) | 48±18 | 53±9 | 54±9 | 60±19 | 0.9 | 0.1 | 0.9 |

Data are means ± SD, \* (p<0.05)
HIIT<sub>85</sub>: Submaximal training group; HIIT<sub>115</sub>: Supramaximal training group; LA, left atrium; PALS: peak atrial longitudinal strain; PACS: peak atrial contraction strain; negative PAS: negative peak atrial strain; RA: right atrium

### Impact of HIIT on ventricle size and function

At baseline, there was no statistical difference in LV size and function between groups (Table 2). LV diastolic and systolic dimensions did not change after 6 weeks of HIIT irrespective of training intensity. Furthermore, LV systolic function parameters (LVEF, LV GLS) and diastolic function parameters (E/A, e’, E/e’) were not influenced by HIIT (Table 3). However, HIIT at 85% maximal aerobic power increased all RV diameters (by 2 ± 4 mm for basal and mid diameters and by 7 ± 9 mm for longitudinal diameters) and significantly reduced RV global longitudinal strain (20 ± 4 vs. 17 ± 3%, p = 0.04; Table 3).

**Table 3.**
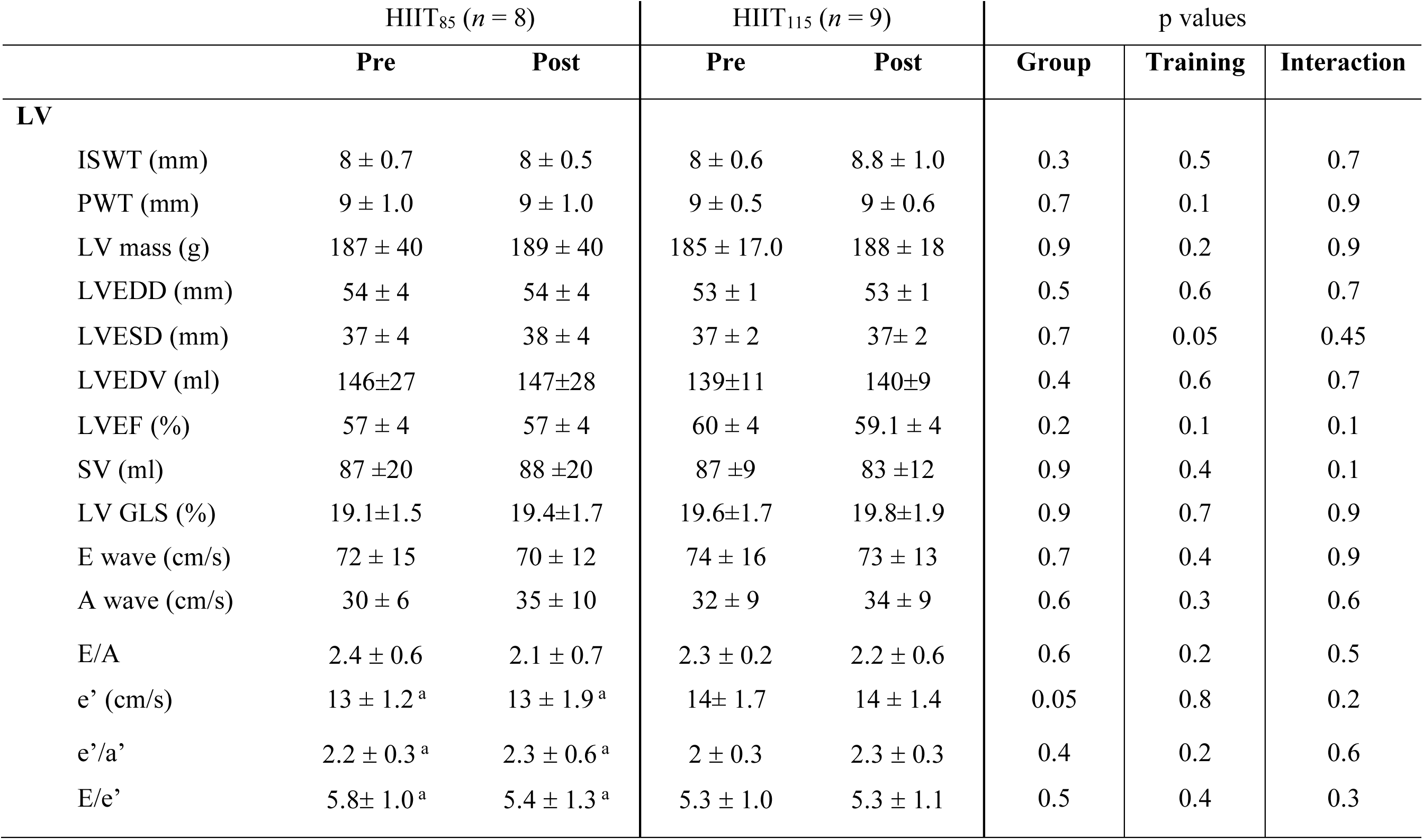
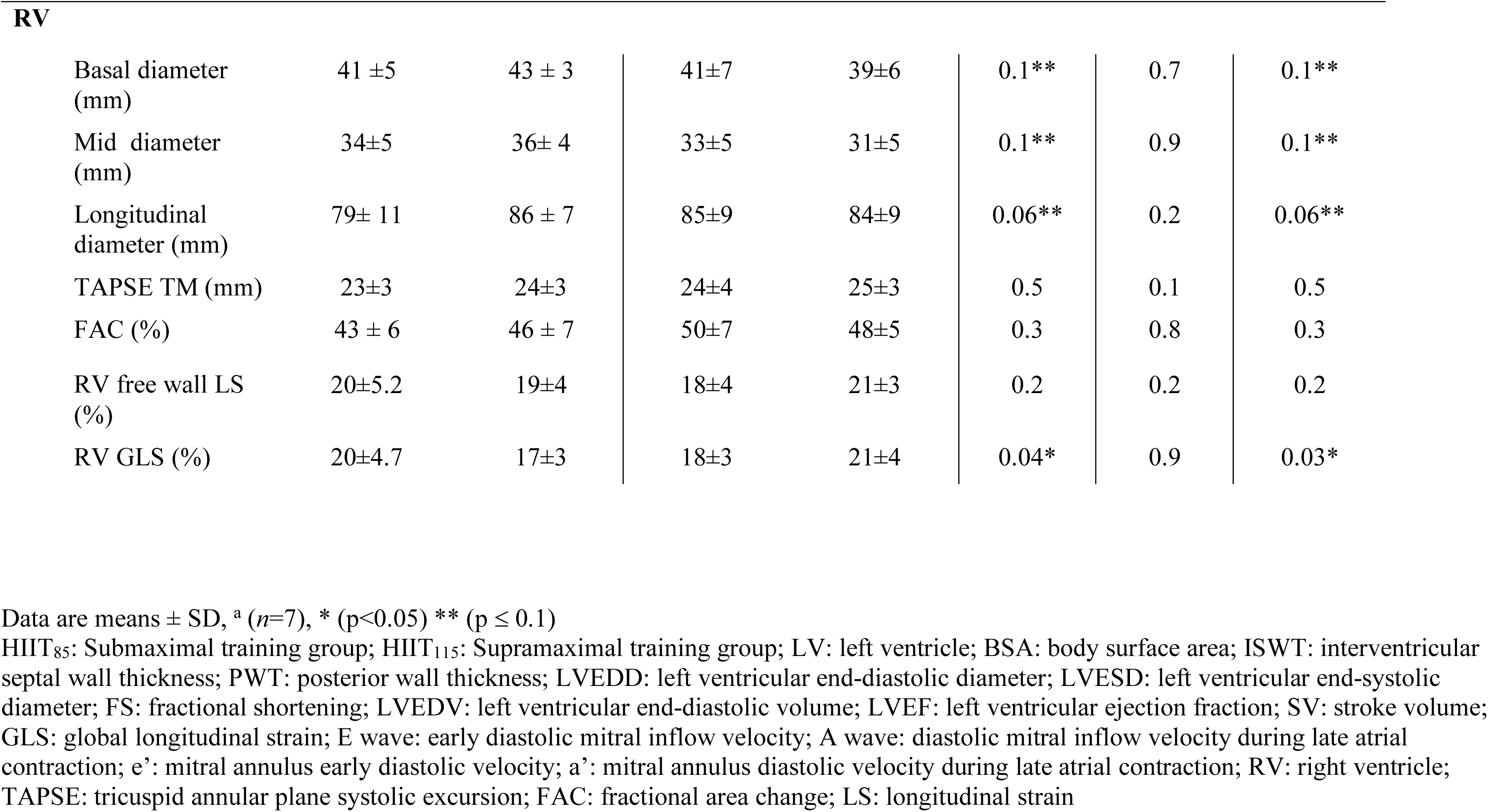
Impact of training on LV and RV size and function in endurance athletes from HIIT_85_ and HIIT_115_ groups.

|  | HIIT<sub>85</sub> (*n* = 8) |  | HIIT<sub>115</sub> (*n* = 9) |  | p values |  |  |
| --- | --- | --- | --- | --- | --- | --- | --- |
|  | Pre | Post | Pre | Post | Group | Training | Interaction |
| **LV** |  |  |  |  |  |  |  |
| ISWT (mm) | 8 ± 0.7 | 8 ± 0.5 | 8 ± 0.6 | 8.8 ± 1.0 | 0.3 | 0.5 | 0.7 |
| PWT (mm) | 9 ± 1.0 | 9 ± 1.0 | 9 ± 0.5 | 9 ± 0.6 | 0.7 | 0.1 | 0.9 |
| LV mass (g) | 187 ± 40 | 189 ± 40 | 185 ± 17.0 | 188 ± 18 | 0.9 | 0.2 | 0.9 |
| LVEDD (mm) | 54 ± 4 | 54 ± 4 | 53 ± 1 | 53 ± 1 | 0.5 | 0.6 | 0.7 |
| LVESD (mm) | 37 ± 4 | 38 ± 4 | 37 ± 2 | 37 ± 2 | 0.7 | 0.05 | 0.45 |
| LVEDV (ml) | 146±27 | 147±28 | 139±11 | 140±9 | 0.4 | 0.6 | 0.7 |
| LVEF (%) | 57 ± 4 | 57 ± 4 | 60 ± 4 | 59.1 ± 4 | 0.2 | 0.1 | 0.1 |
| SV (ml) | 87 ±20 | 88 ±20 | 87 ±9 | 83 ±12 | 0.9 | 0.4 | 0.1 |
| LV GLS (%) | 19.1±1.5 | 19.4±1.7 | 19.6±1.7 | 19.8±1.9 | 0.9 | 0.7 | 0.9 |
| E wave (cm/s) | 72 ± 15 | 70 ± 12 | 74 ± 16 | 73 ± 13 | 0.7 | 0.4 | 0.9 |
| A wave (cm/s) | 30 ± 6 | 35 ± 10 | 32 ± 9 | 34 ± 9 | 0.6 | 0.3 | 0.6 |
| E/A | 2.4 ± 0.6 | 2.1 ± 0.7 | 2.3 ± 0.2 | 2.2 ± 0.6 | 0.6 | 0.2 | 0.5 |
| e' (cm/s) | 13 ± 1.2<sup>a</sup> | 13 ± 1.9<sup>a</sup> | 14 ± 1.7 | 14 ± 1.4 | 0.05 | 0.8 | 0.2 |
| e'/a' | 2.2 ± 0.3<sup>a</sup> | 2.3 ± 0.6<sup>a</sup> | 2 ± 0.3 | 2.3 ± 0.3 | 0.4 | 0.2 | 0.6 |
| E/e' | 5.8 ± 1.0<sup>a</sup> | 5.4 ± 1.3<sup>a</sup> | 5.3 ± 1.0 | 5.3 ± 1.1 | 0.5 | 0.4 | 0.3 |
| **RV** |  |  |  |  |  |  |  |
| Basal diameter (mm) | 41 ±5 | 43 ± 3 | 41±7 | 39±6 | 0.1** | 0.7 | 0.1** |
| Mid diameter (mm) | 34±5 | 36± 4 | 33±5 | 31±5 | 0.1** | 0.9 | 0.1** |
| Longitudinal diameter (mm) | 79± 11 | 86 ± 7 | 85±9 | 84±9 | 0.06** | 0.2 | 0.06** |
| TAPSE TM (mm) | 23±3 | 24±3 | 24±4 | 25±3 | 0.5 | 0.1 | 0.5 |
| FAC (%) | 43 ± 6 | 46 ± 7 | 50±7 | 48±5 | 0.3 | 0.8 | 0.3 |
| RV free wall LS (%) | 20±5.2 | 19±4 | 18±4 | 21±3 | 0.2 | 0.2 | 0.2 |
| RV GLS (%) | 20±4.7 | 17±3 | 18±3 | 21±4 | 0.04* | 0.9 | 0.03* |

Data are means ± SD, <sup>a</sup> (n=7), \* (p<0.05) \*\* (p ≤ 0.1)
HIIT<sub>85</sub>: Submaximal training group; HIIT<sub>115</sub>: Supramaximal training group; LV: left ventricle; BSA: body surface area; ISWT: interventricular septal wall thickness; PWT: posterior wall thickness; LVEDD: left ventricular end-diastolic diameter; LVESD: left ventricular end-systolic diameter; FS: fractional shortening; LVEDV: left ventricular end-diastolic volume; LVEF: left ventricular ejection fraction; SV: stroke volume; GLS: global longitudinal strain; E wave: early diastolic mitral inflow velocity; A wave: diastolic mitral inflow velocity during late atrial contraction; e': mitral annulus early diastolic velocity; a': mitral annulus diastolic velocity during late atrial contraction; RV: right ventricle; TAPSE: tricuspid annular plane systolic excursion; FAC: fractional area change; LS: longitudinal strain

## DISCUSSION

The main findings of this study are: 1) a significant decrease in resting heart rate, ambulatory SBP, MAP and PP, 2) a significant increase in LA volumes after six weeks of HIIT and, 3) a significant decrease in RV systolic function evaluated by longitudinal strain in endurance-trained males who underwent a submaximal HIIT protocol (HIIT_85_) associated with increased RV dimensions. To the best of our knowledge, this is the first study to report significant structural and functional cardiac changes in endurance-trained individuals after only 6 weeks of well-defined HIIT protocols to exhaustion.

### HIIT and blood pressure

Few studies have evaluated the impact of HIIT on BP measured by ambulatory BP monitoring, a measurement that is more specific for systemic arterial hypertension diagnosis (Ciolac *et al*., 2010; Santos *et al*., 2016; Whelton *et al*., 2018). Our results represent clinically positive findings for endurance-trained participants since systemic arterial hypertension is the most common cardiovascular disease in these individuals (Leddy & Izzo, 2009) and it is often masked (Trachsel *et al*., 2015). In our study, our three participants that were considered mildly hypertensive at baseline normalized their ambulatory BP after HIIT.

Continuous moderate-intensity exercise training (CMT), that can be sustained for 30 min or more, has traditionally been recommended for systemic arterial hypertension prevention and treatment (Haskell *et al*., 2007). However, the literature related to the impact of intensity training on BP in normotensive and hypertensive participants remains equivocal. Indeed, several studies reported that CMT and HIIT were equally effective at reducing ambulatory BP (Ciolac *et al*., 2009; Guimaraes *et al*., 2010), individuals with highest baseline values experiencing the greatest BP reductions (Pescatello & Kulikowich, 2001). However, HIIT was found to be superior to CMT at improving endothelial dysfunction, sympathetic nerve activity and arterial stiffness in hypertensive patients and normotensive individuals at high risk for hypertension (Tjonna *et al*., 2008; Ciolac *et al*., 2010; Guimaraes *et al*., 2010; Rakobowchuk *et al*., 2013). The decrease in ambulatory SBP and MAP observed in the current study may be explained by diminished arterial stiffness after HIIT, which would support the theory that HIIT provides more peripheral adaptations (Macpherson *et al*., 2011). Improvements in endothelial function and sympathetic nerve activity of arterial smooth muscle cells are possible mechanisms involved in arterial stiffness reduction induced by exercise training (Zieman *et al*., 2005).

### Exercise-induced cardiac remodeling and its possible consequences

Highly trained individuals develop cardiac adaptations including enlarged LV and RV volumes, increased LV wall thickness and cardiac mass, and increased LA size (Pelliccia *et al*., 1991; Pelliccia *et al*., 2005). These structural alterations, together with a preserved LVEF as well as a LV diastolic function, have been considered typical findings of the “athlete’s heart” (Spirito *et al*., 1994; Pluim *et al*., 2000; Maron & Pelliccia, 2006). Accumulating data suggest that some of the remodeling that occurs in endurance athletes may not be entirely benign (Ector *et al*., 2007; Mohlenkamp *et al*., 2008; Breuckmann *et al*., 2009; Trivax *et al*., 2010).

Our study reported a significant increase in LA volumes after a short 6-week HIIT protocol to exhaustion and a significant RV dysfunction in participants of the submaximal HIIT group associated with an increase in RV dimensions and no effect on LV structure and function. Of note, our endurance-trained participants had no features of the athlete’s heart at baseline. Several studies reported LA enlargement after a training period. D’Ascenzi et al. observed in a population of adolescent soccer players an increase in LA volumes, which occurred after 4 months of intensive training with further increment after 8 months (D’Ascenzi *et al*., 2012). Baggish et al. found similar results after 90 days of intense training in a group of endurance athletes, but not in a group of strength-trained athletes (Baggish *et al*., 2008). Atrial enlargement observed in athletes represents an adaptive mechanism to the increased volume overload induced by training. This process is dynamic and reversible. However, the extent of LA dimensional remodeling in highly trained athletes may be relevant and absolute LA size can overlap atrial dilation observed in patients with cardiac disease and can be observed in the growing hearts of young athletes (D’Ascenzi *et al*., 2016).

In regards to the influence of exercise training on right cardiac chambers, acute changes were observed in previous studies. Trivax et al. found acute RA and RV dilation with concomitant RV dysfunction in 25 runners immediately after a marathon whereas high values of cardiac troponin I and B-type natriuretic peptide were also observed (Trivax *et al*., 2010). In a group of 40 athletes, La Gerche et al. reported acute RV dilation and dysfunction immediately after an endurance race (3-11 hours) with elevated cardiac troponin and B-natriuretic peptide blood levels but no changes in LV volumes and function. In that study, RV dysfunction (mostly recovered after one week) decreased with longer race duration, and myocardial scarring was detected by focal gadolinium enhancement localized to the interventricular septum on cardiac magnetic resonance imaging in five athletes who had greater exercise exposure and lower RV ejection fraction (La Gerche *et al*., 2012). Of interest, in our study, only participants who underwent a submaximal HIIT protocol had increased RV dimensions and RV dysfunction. These endurance-trained males had lower intensity exercise training but higher exercise volume (1.4-fold more than the HIIT_115_ group). All these results suggest that intense endurance exercise induces acute RV dilation and RV dysfunction while sparing LV function. The RV is known to be very sensitive to volume overload because of its thin wall, recurrent stretch of the chamber and reestablishment of the chamber geometry (O’Keefe *et al*., 2012). In some individuals, the RV may be prone to the development of chronic structural changes including chronic dilation of the RV and RA, scattered areas of myocardial injury with subsequent fibrotic scarring and subsequent RV dysfunction (O’Keefe *et al*., 2012). Evidence of patchy myocardial scarring assessed by magnetic resonance imaging has also been observed in 12% of healthy male runners aged from 50 to 72 years old who had completed at least 5 marathons during the past three years; a rate 3-fold higher than that in age-matched control subjects (Breuckmann *et al*., 2009). Moreover, complex ventricular ectopy, including ventricular tachycardia and sudden cardiac death have been reported in very fit individuals (Kim *et al*., 2012). The ventricular arrhythmias typically originate from a mildly dysfunctional RV and/or the interventricular septum. The patchy myocardial fibrosis that may develop as a reparative response to damaged myocardium can favor reentry, which is well established as a substrate for arrhythmia (Biffi *et al*., 2002; Heidbuchel *et al*., 2003; Ector *et al*., 2007; Burstein & Nattel, 2008; Oxborough *et al*., 2010).

These observations have led to speculation about the existence of a syndrome of exercise-induced arrhythmogenic RV cardiomyopathy, that shares some features with the familial RV disease but is caused by chronic high-level endurance exercise training rather than genetic predisposition (Breuckmann *et al*., 2009; Trivax *et al*., 2010). Similarly, significant associations have been reported between LA dilation and prevalence of atrial fibrillation among long-term endurance athletes (Molina *et al*., 2008; Grimsmo *et al*., 2010) with as much as a 5-fold increase in this prevalence with long-term sustained vigorous aerobic training (Mont *et al*., 2002; Neilan *et al*., 2006; Aizer *et al*., 2009; Mont *et al*., 2009). Long-term structural changes in the LA, LA fibrosis with increased LA stiffness and increased vagal tone related to high-intensity exercise are the main hypothesized mechanisms, the myocardial fibrosis creating a substrate for atrial arrhythmias (Pluim *et al*., 2000; Maron & Pelliccia, 2006; Ector *et al*., 2007; Nassenstein *et al*., 2009).

### Clinical Implications

Exercise has substantial health benefits with pleiomorphic, cardiovascular, metabolic, psychological, anti-neoplastic actions resulting in improved quality of life and longevity as endurance athletes on average being healthier and living longer than non-athletes (Sarna *et al*., 1993). Our study results found a significant reduction in ambulatory BP in a group of endurance-trained males after only 6 weeks of HIIT to exhaustion enhancing the positive benefits of intense exercise training in the management of systemic arterial hypertension, one of the major risk factors of cardiac morbidity and mortality worldwide (Kokubo & Matsumoto, 2017).

Moreover, as low adherence in exercise training is often associated with lack of time (Gillen & Gibala, 2014), HIIT may be a cardiovascular exercise strategy to promote greater adherence as it covers relatively short training sessions (20-25min), comprising high-intensity exercise intervals interspersed with rest or active recovery, allowing patients to have complete periods at high intensity. Moreover, the periods of high-intensity exercise stimulate more intense cardiovascular and muscular adaptations in patients with heart disease, which favors mitochondrial function and calcium reabsorption rate into the sarcoplasmic reticulum, reducing skeletal muscle fatigue and increasing exercise capacity (Wisloff *et al*., 2007; Ito *et al*., 2016). However, our study also highlights, in agreement with numerous previous study data, the fact that endurance-trained individuals are likely to develop LA and RV volumes increase as well as subclinical RV dysfunction. These changes, which are observed soon after intense periods of training, may become chronic in highly trained athletes and in athletes with chronic excessive endurance training such as those who train/compete in extreme endurance training (O’Keefe *et al*., 2012). Moreover, exercise-induced increase in atrial and RV volumes have been associated with myocardial fibrosis and with incidence of atrial fibrillation and ventricular arrhythmias. Further studies are obviously needed to understand the potential mechanisms underlying these associations and to determine the long-term clinical significance of these cardiac changes. However, the type, intensity and amount of exercise training appear to influence the risk of adverse cardiovascular outcomes.

It is important to highlight that our participants exercised until exhaustion during each HIIT session of their exercise protocol. Similar protocols have been used to study the effects of overtraining in endurance athletes inducing stroke volume and cardiac output decreases during exercise and increased resting arterial stiffness (Cote *et al*., 2013; Coates *et al*., 2018). Therefore, one can wonder whether it is possible to determine a safe upper range dose of exercise training above which the adverse cardiac effects of sustained intense training may outweigh its benefits in order to help elaborating safer and more adapted training programs for athletes especially in highly trained, and veteran endurance athletes.

### Limitations

The present study is a prospective study with ambulatory BP measurements and echocardiographic exams performed at baseline and 48h to 96h after ending our HIIT protocols. No evaluation has been done later to determine if observed changes were transient. No women were included in the group study impacting the generalizability of our observations. The apparent lack of significant changes in LA function evaluated by strain after HIIT protocols may be a type II error because of the small sample size. Nonetheless, this limitation does not affect the validity of the main results of the study, which are a significant increase in LA volumes in all participants after only 6 weeks of HIIT and a significant decrease in subclinical RV function in the HIIT_85_ group.

## CONCLUSION

This study reports that six weeks of both submaximal and supramaximal interval training performed to exhaustion thrice weekly significantly reduce ambulatory BP and increase LA volumes with no effect on LV structure and function in endurance-trained individuals. In the submaximal training group, RV dimensions increased with a deterioration of RV function assessed by strain imaging. The positive impact of HIIT on ambulatory BP, its capacity to improve physical performance in already fit participants and its better adherence due to shorter training sessions makes HIIT a promising exercise strategy for endurance athletes and for individuals with cardiometabolic diseases. In light of observed atrial and right ventricle changes in structure and function after intense exercise training to exhaustion and the possible association of these changes with incidence of atrial and ventricular arrhythmias in the future, further investigation is warranted to identify the exercise threshold for potential “toxicity” of intense exercise training for at-risk individuals and ideal exercise training regimens conferring optimal cardiovascular protection and adapted endurance training for athletes.

## ADDITIONNAL INFORMATION

### Author contributions

MP and PB conceived the study and designed the experiment. HM, OL, MP, SI, LL, AD, EB and PB contributed to data collection and/or analysis. HM drafted the manuscript. OL, MP, SI, LL, AD, PPo, EB and PPi contributed to critically revising of this manuscript. All authors approved the final version.

### Funding

The present study was funded by the Ministère de l’Éducation, du Loisir et du Sport du Québec, and the Foundation of the Institut universitaire de cardiologie et de pneumologie de Québec. Myriam Paquette received a scholarship from the Canadian Institutes of Health Research. SI was supported by a doctoral training scholarship from the Fonds de recherche du Québec – Santé. LL is supported by a doctoral training scholarship from the Société Québécoise d’hypertension artérielle. Paul Poirier is a senior clinical-scientist of the Fonds de recherche du Québec – Santé.

## Acknowledgements

We express our gratitude to the participants for their enthusiastic participation during the vigorous training protocol and for their availability to all the testing and training sessions. We also want to thank Louis-Charles B. Lacroix and Andrée-Anne Clément for their assistance in the supervision of training sessions, Sophie Castonguay-Paradis for her help in data collection and Lise Renaud for their work and echocardiography data review. We also want to thank Tacx Canada and most specifically Cycle Lambert, for lending us Tacx Bushido home trainers.

## REFERENCES

Aizer A, Gaziano JM, Cook NR, Manson JE, Buring JE & Albert CM. (2009). Relation of vigorous exercise to risk of atrial fibrillation. Am J Cardiol 103, 1572–1577.

Ancona R, Comenale Pinto S, Caso P, D’Andrea A, Di Salvo G, Arenga F, Coppola MG, Sellitto V, Macrino M & Calabro R. (2014). Left atrium by echocardiography in clinical practice: from conventional methods to new echocardiographic techniques. ScientificWorldJournal 2014, 451042.

Baggish AL, Wang F, Weiner RB, Elinoff JM, Tournoux F, Boland A, Picard MH, Hutter AM, Jr. & Wood MJ. (2008). Training-specific changes in cardiac structure and function: a prospective and longitudinal assessment of competitive athletes. J Appl Physiol 104, 1121–1128.

Batacan RB, Jr., Duncan MJ, Dalbo VJ, Tucker PS & Fenning AS. (2017). Effects of high-intensity interval training on cardiometabolic health: a systematic review and meta-analysis of intervention studies. Br J Sports Med 51, 494–503.

Berge HM, Isern CB & Berge E. (2015). Blood pressure and hypertension in athletes: a systematic review. Br J Sports Med.

Biffi A, Pelliccia A, Verdile L, Fernando F, Spataro A, Caselli S, Santini M & Maron BJ. (2002). Long-term clinical significance of frequent and complex ventricular tachyarrhythmias in trained athletes. J Am Coll Cardiol 40, 446–452.

Breuckmann F, Mohlenkamp S, Nassenstein K, Lehmann N, Ladd S, Schmermund A, Sievers B, Schlosser T, Jockel KH, Heusch G, Erbel R & Barkhausen J. (2009). Myocardial late gadolinium enhancement: prevalence, pattern, and prognostic relevance in marathon runners. Radiology 251, 50–57.

Buchheit M & Laursen PB. (2013). High-intensity interval training, solutions to the programming puzzle: Part I: cardiopulmonary emphasis. Sports Med 43, 313–338.

Burstein B & Nattel S. (2008). Atrial fibrosis: mechanisms and clinical relevance in atrial fibrillation. J Am Coll Cardiol 51, 802–809.

Chobanian AV. (2007). Clinical practice. Isolated systolic hypertension in the elderly. N Engl J Med 357, 789–796.

Ciolac EG, Bocchi EA, Bortolotto LA, Carvalho VO, Greve JM & Guimaraes GV. (2010). Effects of high-intensity aerobic interval training vs. moderate exercise on hemodynamic, metabolic and neuro-humoral abnormalities of young normotensive women at high familial risk for hypertension. Hypertens Res 33, 836–843.

Ciolac EG, Guimaraes GV, VM DA, Bortolotto LA, Doria EL & Bocchi EA. (2009). Acute effects of continuous and interval aerobic exercise on 24-h ambulatory blood pressure in long-term treated hypertensive patients. Int J Cardiol 133, 381–387.

Coates AM, Millar PJ & Burr JF. (2018). Blunted Cardiac Output from Overtraining Is Related to Increased Arterial Stiffness. Med Sci Sports Exerc 50, 2459–2464.

Cornelissen VA & Smart NA. (2013). Exercise training for blood pressure: a systematic review and meta-analysis. Journal of the American Heart Association 2, e004473.

Cote AT, Bredin SS, Phillips AA, Koehle MS, Glier MB, Devlin AM & Warburton DE. (2013). Left ventricular mechanics and arterial-ventricular coupling following high-intensity interval exercise. J Appl Physiol (1985) 115, 1705–1713.

D’Ascenzi F, Cameli M, Lisi M, Zaca V, Natali B, Malandrino A, Benincasa S, Catanese S, Causarano A & Mondillo S. (2012). Left atrial remodelling in competitive adolescent soccer players. Int J Sports Med 33, 795–801.

D’Ascenzi F, Pelliccia A, Natali BM, Cameli M, Lisi M, Focardi M, Padeletti M, Palmitesta P, Corrado D, Bonifazi M, Mondillo S & Henein M. (2015). Training-induced dynamic changes in left atrial reservoir, conduit, and active volumes in professional soccer players. Eur J Appl Physiol 115, 1715–1723.

D’Ascenzi F, Solari M, Anselmi F, Maffei S, Focardi M, Bonifazi M, Mondillo S & Henein M. (2016). Atrial chamber remodelling in healthy pre-adolescent athletes engaged in endurance sports: A study with a longitudinal design. The CHILD study. Int J Cardiol 223, 325–330.

Ector J, Ganame J, van der Merwe N, Adriaenssens B, Pison L, Willems R, Gewillig M & Heidbuchel H. (2007). Reduced right ventricular ejection fraction in endurance athletes presenting with ventricular arrhythmias: a quantitative angiographic assessment. Eur Heart J 28, 345–353.

Foster C. (1998). Monitoring training in athletes with reference to overtraining syndrome. Med Sci Sports Exerc 30, 1164–1168.

Gillen JB & Gibala MJ. (2014). Is high-intensity interval training a time-efficient exercise strategy to improve health and fitness? Appl Physiol Nutr Metab 39, 409–412.

Grimsmo J, Grundvold I, Maehlum S & Arnesen H. (2010). High prevalence of atrial fibrillation in long-term endurance cross-country skiers: echocardiographic findings and possible predictors--a 28-30 years follow-up study. Eur J Cardiovasc Prev Rehabil 17, 100–105.

Guimaraes GV, Ciolac EG, Carvalho VO, D’Avila VM, Bortolotto LA & Bocchi EA. (2010). Effects of continuous vs. interval exercise training on blood pressure and arterial stiffness in treated hypertension. Hypertens Res 33, 627–632.

Haskell WL, Lee IM, Pate RR, Powell KE, Blair SN, Franklin BA, Macera CA, Heath GW, Thompson PD, Bauman A, American College of Sports M & American Heart A. (2007). Physical activity and public health: updated recommendation for adults from the American College of Sports Medicine and the American Heart Association. Circulation 116, 1081–1093.

Heidbuchel H, Hoogsteen J, Fagard R, Vanhees L, Ector H, Willems R & Van Lierde J. (2003). High prevalence of right ventricular involvement in endurance athletes with ventricular arrhythmias. Role of an electrophysiologic study in risk stratification. Eur Heart J 24, 1473–1480.

Ito S, Mizoguchi T & Saeki T. (2016). Review of High-intensity Interval Training in Cardiac Rehabilitation. Intern Med 55, 2329–2336.

Kim JH, Malhotra R, Chiampas G, d’Hemecourt P, Troyanos C, Cianca J, Smith RN, Wang TJ, Roberts WO, Thompson PD, Baggish AL & Race Associated Cardiac Arrest Event Registry Study G. (2012). Cardiac arrest during long-distance running races. N Engl J Med 366, 130–140.

Kokubo Y & Matsumoto C. (2017). Hypertension Is a Risk Factor for Several Types of Heart Disease: Review of Prospective Studies. Adv Exp Med Biol 956, 419–426.

La Gerche A, Burns AT, Mooney DJ, Inder WJ, Taylor AJ, Bogaert J, Macisaac AI, Heidbuchel H & Prior DL. (2012). Exercise-induced right ventricular dysfunction and structural remodelling in endurance athletes. Eur Heart J 33, 998–1006.

Lang RM, Badano LP, Mor-Avi V, Afilalo J, Armstrong A, Ernande L, Flachskampf FA, Foster E, Goldstein SA, Kuznetsova T, Lancellotti P, Muraru D, Picard MH, Rietzschel ER, Rudski L, Spencer KT, Tsang W & Voigt JU. (2015). Recommendations for cardiac chamber quantification by echocardiography in adults: an update from the American Society of Echocardiography and the European Association of Cardiovascular Imaging. J Am Soc Echocardiogr 28, 1–39 e14.

Leddy JJ & Izzo J. (2009). Hypertension in athletes. J Clin Hypertens 11, 226–233.

Machino-Ohtsuka T, Seo Y, Tada H, Ishizu T, Machino T, Yamasaki H, Igarashi M, Xu D, Sekiguchi Y & Aonuma K. (2011). Left atrial stiffness relates to left ventricular diastolic dysfunction and recurrence after pulmonary vein isolation for atrial fibrillation. J Cardiovasc Electrophysiol 22, 999–1006.

Macpherson RE, Hazell TJ, Olver TD, Paterson DH & Lemon PW. (2011). Run sprint interval training improves aerobic performance but not maximal cardiac output. Med Sci Sports Exerc 43, 115–122.

Maron BJ & Pelliccia A. (2006). The heart of trained athletes: cardiac remodeling and the risks of sports, including sudden death. Circulation 114, 1633–1644.

Mohlenkamp S, Lehmann N, Breuckmann F, Brocker-Preuss M, Nassenstein K, Halle M, Budde T, Mann K, Barkhausen J, Heusch G, Jockel KH, Erbel R, Marathon Study I & Heinz Nixdorf Recall Study I. (2008). Running: the risk of coronary events: Prevalence and prognostic relevance of coronary atherosclerosis in marathon runners. Eur Heart J 29, 1903–1910.

Molina L, Mont L, Marrugat J, Berruezo A, Brugada J, Bruguera J, Rebato C & Elosua R. (2008). Long-term endurance sport practice increases the incidence of lone atrial fibrillation in men: a follow-up study. Europace 10, 618–623.

Mont L, Elosua R & Brugada J. (2009). Endurance sport practice as a risk factor for atrial fibrillation and atrial flutter. Europace 11, 11–17.

Mont L, Sambola A, Brugada J, Vacca M, Marrugat J, Elosua R, Pare C, Azqueta M & Sanz G. (2002). Long-lasting sport practice and lone atrial fibrillation. Eur Heart J 23, 477–482.

Nassenstein K, Breuckmann F, Lehmann N, Schmermund A, Hunold P, Broecker-Preuss M, Sandner TA, Halle M, Mann K, Jockel KH, Heusch G, Budde T, Erbel R, Barkhausen J & Mohlenkamp S. (2009). Left ventricular volumes and mass in marathon runners and their association with cardiovascular risk factors. Int J Cardiovasc Imaging 25, 71–79.

Neilan TG, Januzzi JL, Lee-Lewandrowski E, Ton-Nu TT, Yoerger DM, Jassal DS, Lewandrowski KB, Siegel AJ, Marshall JE, Douglas PS, Lawlor D, Picard MH & Wood MJ. (2006). Myocardial injury and ventricular dysfunction related to training levels among nonelite participants in the Boston marathon. Circulation 114, 2325–2333.

O’Keefe JH, Patil HR, Lavie CJ, Magalski A, Vogel RA & McCullough PA. (2012). Potential adverse cardiovascular effects from excessive endurance exercise. Mayo Clin Proc 87, 587–595.

Oxborough D, Birch K, Shave R & George K. (2010). “Exercise-induced cardiac fatigue”--a review of the echocardiographic literature. Echocardiography 27, 1130–1140.

Paquette M, Le Blanc O, Lucas SJ, Thibault G, Bailey DM & Brassard P. (2017). Effects of submaximal and supramaximal interval training on determinants of endurance performance in endurance athletes. Scand J Med Sci Sports 27, 318–326.

Pelliccia A, Maron BJ, Di Paolo FM, Biffi A, Quattrini FM, Pisicchio C, Roselli A, Caselli S & Culasso F. (2005). Prevalence and clinical significance of left atrial remodeling in competitive athletes. J Am Coll Cardiol 46, 690–696.

Pelliccia A, Maron BJ, Spataro A, Proschan MA & Spirito P. (1991). The upper limit of physiologic cardiac hypertrophy in highly trained elite athletes. N Engl J Med 324, 295–301.

Pescatello LS & Kulikowich JM. (2001). The aftereffects of dynamic exercise on ambulatory blood pressure. Med Sci Sports Exerc 33, 1855–1861.

Pluim BM, Zwinderman AH, van der Laarse A & van der Wall EE. (2000). The athlete’s heart. A meta-analysis of cardiac structure and function. Circulation 101, 336–344.

Rakobowchuk M, Harris E, Taylor A, Cubbon RM & Birch KM. (2013). Moderate and heavy metabolic stress interval training improve arterial stiffness and heart rate dynamics in humans. Eur J Appl Physiol 113, 839–849.

Ramos JS, Dalleck LC, Tjonna AE, Beetham KS & Coombes JS. (2015). The impact of high-intensity interval training versus moderate-intensity continuous training on vascular function: a systematic review and meta-analysis. Sports Med 45, 679–692.

Santos LP, Moraes RS, Vieira PJ, Ash GI, Waclawovsky G, Pescatello LS & Umpierre D. (2016). Effects of aerobic exercise intensity on ambulatory blood pressure and vascular responses in resistant hypertension: a crossover trial. J Hypertens 34, 1317–1324.

Sarna S, Sahi T, Koskenvuo M & Kaprio J. (1993). Increased life expectancy of world class male athletes. Med Sci Sports Exerc 25, 237–244.

Spirito P, Pelliccia A, Proschan MA, Granata M, Spataro A, Bellone P, Caselli G, Biffi A, Vecchio C & Maron BJ. (1994). Morphology of the “athlete’s heart” assessed by echocardiography in 947 elite athletes representing 27 sports. Am J Cardiol 74, 802–806.

Tjonna AE, Lee SJ, Rognmo O, Stolen TO, Bye A, Haram PM, Loennechen JP, Al-Share QY, Skogvoll E, Slordahl SA, Kemi OJ, Najjar SM & Wisloff U. (2008). Aerobic interval training versus continuous moderate exercise as a treatment for the metabolic syndrome: a pilot study. Circulation 118, 346–354.

Trachsel LD, Carlen F, Brugger N, Seiler C & Wilhelm M. (2015). Masked hypertension and cardiac remodeling in middle-aged endurance athletes. J Hypertens 33, 1276–1283.

Trivax JE, Franklin BA, Goldstein JA, Chinnaiyan KM, Gallagher MJ, deJong AT, Colar JM, Haines DE & McCullough PA. (2010). Acute cardiac effects of marathon running. J Appl Physiol 108, 1148–1153.

Westgarth-Taylor C, Hawley JA, Rickard S, Myburgh KH, Noakes TD & Dennis SC. (1997). Metabolic and performance adaptations to interval training in endurance-trained cyclists. Eur J Appl Physiol Occup Physiol 75, 298–304.

Weston M, Taylor KL, Batterham AM & Hopkins WG. (2014). Effects of low-volume high-intensity interval training (HIT) on fitness in adults: a meta-analysis of controlled and non-controlled trials. Sports Med 44, 1005–1017.

Whelton PK, Carey RM, Aronow WS, Casey DE, Jr., Collins KJ, Dennison Himmelfarb C, DePalma SM, Gidding S, Jamerson KA, Jones DW, MacLaughlin EJ, Muntner P, Ovbiagele B, Smith SC, Jr., Spencer CC, Stafford RS, Taler SJ, Thomas RJ, Williams KA, Sr., Williamson JD & Wright JT, Jr. (2018). 2017 Guideline for the Prevention, Detection, Evaluation, and Management of High Blood Pressure in Adults: A Report of the American College of Cardiology/American Heart Association Task Force on Clinical Practice Guidelines. Hypertension 71, e13–e115.

Wisloff U, Stoylen A, Loennechen JP, Bruvold M, Rognmo O, Haram PM, Tjonna AE, Helgerud J, Slordahl SA, Lee SJ, Videm V, Bye A, Smith GL, Najjar SM, Ellingsen O & Skjaerpe T. (2007). Superior cardiovascular effect of aerobic interval training versus moderate continuous training in heart failure patients: a randomized study. Circulation 115, 3086–3094.

Zieman SJ, Melenovsky V & Kass DA. (2005). Mechanisms, pathophysiology, and therapy of arterial stiffness. Arterioscler Thromb Vasc Biol 25, 932–943.

